# Myeloid Cell Glucocorticoid, Not Mineralocorticoid Receptor Signaling, Contributes to Salt-Sensitive Hypertension in Humans via Cortisol

**DOI:** 10.1101/2024.06.10.598374

**Authors:** Claude F. Albritton, Mert Demirci, Kit Neikirk, Lale A. Ertuglu, Jeanne A Ishimwe, Ashley L Mutchler, Quanhu Sheng, Cheryl L Laffer, Celestine N. Wanjalla, Taseer Ahmed, Alexandria Porcia Haynes, Mohammad Saleem, Heather K. Beasley, Andrea G. Marshall, Zer Vue, Alp T Ikizler, Thomas R. Kleyman, Valentina Kon, Antentor Hinton, Annet Kirabo

## Abstract

**BACKGROUND:** Salt sensitivity of blood pressure (SSBP) is an independent risk factor for cardiovascular morbidity and mortality, yet the etiology is poorly understood. We previously found that serum/glucocorticoid-regulated kinase 1 (SGK1) and epoxyeicosatrienoic acids (EETs) regulate epithelial sodium channel (ENaC)-dependent sodium entry into monocyte-derived antigen-presenting cells (APCs) and activation of NADPH oxidase, leading to the formation of isolevuglandins (IsoLGs) in SSBP. Whereas aldosterone via the mineralocorticoid receptor (MR) activates SGK1 leading to hypertension, our past findings indicate that levels of plasma aldosterone do not correlate with SSBP, and there is little to no MR expression in APCs. Thus, we hypothesized that cortisol acting via the glucocorticoid receptor (GR), not the MR in APCs mediates SGK1 actions to induce SSBP.

**METHODS:** We performed cellular indexing of transcriptomes and epitopes by sequencing (CITE-Seq) analysis on peripheral blood mononuclear cells of humans rigorously phenotyped for SSBP using an inpatient salt loading/depletion protocol to determine expression of MR, GR, and SGK1 in immune cells. In additional experiments, we performed bulk transcriptomic analysis on isolated human monocytes following *in vitro* treatment with high salt from a separate cohort. We then measured urine and plasma cortisol, cortisone, renin, and aldosterone. Subsequently, we measured the association of these hormones with changes in systolic, diastolic, mean arterial pressure and pulse pressure as well as immune cell activation via IsoLG formation.

**RESULTS:** We found that myeloid APCs predominantly express the GR and SGK1 with no expression of the MR. Expression of the GR in APCs increased after salt loading and decreased with salt depletion in salt-sensitive but not salt-resistant people and was associated with increased expression of *SGK1*. Moreover, we found that plasma and urine cortisol/cortisone but not aldosterone/renin correlated with SSBP and APCs activation via IsoLGs. We also found that cortisol negatively correlates with EETs.

**CONCLUSION:** Our findings suggest that renal cortisol signaling via the GR but not the MR in APCs contributes to SSBP via cortisol. Urine and plasma cortisol may provide an important currently unavailable feasible diagnostic tool for SSBP. Moreover, cortisol-GR-SGK1-ENaC signaling pathway may provide treatment options for SSBP.

**Novelty and Relevance:** *What Is New?:*

- Although salt sensitivity is a major risk factor for cardiovascular morbidity and mortality, the mechanisms underlying the salt sensitivity of blood pressure (SSBP) are poorly understood.
- High salt modifies glucocorticoid-receptor expression in antigen-presenting cells (APCs), suggesting a critical role of glucocorticoids in SSBP.
- Elevated glucocorticoid receptor (GR) expression compared to mineralocorticoid receptor (MR) expression in APCs provides evidence for a GR-dependent pathway to SSBP. Isolevuglandins (IsoLGs) increased in APCs *in vitro* after hydrocortisone treatment compared to aldosterone treatment, indicating that cortisol was the predominant driver of IsoLG production in these cells.
- Our studies suggest a mechanism for *SGK1* expression through GR activation by cortisol that differs from the currently accepted mechanism for SSBP pathogenesis.

*What Is Relevant?:*

- Although aldosterone has been used to study SSBP, there has been no consideration of cortisol as a major driver of the condition.
- Understanding alternative inflammatory pathways that affect SSBP may provide insights into the mechanism of SSBP and suggest a range of therapeutic targets.
- Our studies may provide a practical approach to understanding and treating salt-sensitive hypertension.

*Clinical/Pathophysiological Implications?:*

- Our findings firmly support a GR-dependent signaling pathway for activating SSBP via *SGK1* expression. A cortisol-driven mechanism could provide a practical approach for targeted treatments for salt-sensitive hypertension. Moreover, it could pave the way for a diagnostic approach.

## Introduction

Globally, dietary salt consumption has increased with the evolution of processed foods, leading to a greater risk of hypertension and cardiovascular disease.^1–3^ Salt sensitivity of blood pressure (SSBP) refers to the heterogeneity in blood pressure responses according to dietary salt intake.^1^ Individuals who respond to high dietary salt intake with a greater blood pressure increase in an acute manner are classified as salt-sensitive (SS)—a condition that has commonly been linked to genetic abnormalities^4^—and face a higher risk of cardiovascular events and deaths due to excessive salt intake compared to salt-resistant (SR) individuals.^1–5^ The etiology of SSBP is not clearly understood, yet mechanistic investigation is hampered by the absence of feasible diagnostic screening tools.

Isolevuglandins (IsoLGs) are highly reactive products of lipid oxidation that form covalent bonds with lysine residues, leading to post-translational protein modifications.^6^ We found that IsoLGs accumulate and act as neoantigens in antigen-presenting cells (APCs) to activate T cells and promote hypertension.^7,8^ Our studies indicate that dietary sodium (Na^+^) is a potent stimulus for IsoLG-adduct formation in APCs via activation of the NADPH oxidase.^9^ Mice lacking NADPH oxidase do not form IsoLG adducts, and pharmacological scavenging of IsoLGs prevents dendritic cell (DC) activation, hypertension, and end-organ damage.^6,9^ Thus, therapeutic strategies to reduce Na^+^ intake or signaling may reduce inflammation and hypertension.^10^

The epithelial sodium channel (ENaC), which is implicated in the pathogenesis of SSBP^11–13^, has been studied extensively in the kidney, where it regulates electrolyte and volume balance. Our studies indicate that dendritic cell (DC) entry of extracellular Na^+^ requires ENaC, which is then exchanged for Ca^2+^ via the Na^+^/Ca^2+^ exchanger with the resultant activation of protein kinase C (PKC). PKC phosphorylates the p47 subunit of NADPH oxidase leading to formation of superoxide and downstream immunogenic IsoLG-adducts in APCs.^6^ These APCs activate T cells and inflammatory cytokines which induce vascular and renal Na^+^ transporter dysfunction and increase in blood pressure.^8^ We found that the expression of ENaC in APCs in response to high salt is regulated by both epoxyeicosatrienoic acids (EETs) and serum/glucocorticoid regulated kinase 1 (SGK1).^14^ Moreover, we found that SGK1 plays a role in SSBP as its expression in APCs parallel blood pressure changes during salt loading/depletion in SS people.^15^ However, the upstream signaling mechanisms leading to the expression of APC SGK1 in SSBP are unknown.

Medical conditions that result in excessive cortisol have been associated with hypertension and are associated with defects in 11β-hydroxysteroid dehydrogenases (11β-HSDs)^16^. Impairment in 11β-HSD increases cortisol in peripheral tissues, further exacerbated by high salt intake.^16–18^ Excess cortisol and 11-deoxycortisol, as well as elevated cortisol-to-cortisone ratios, have been implicated in hypertension^19^. Although aldosterone typically binds to the mineralocorticoid receptor (MR) to increase membrane expression of ENaC via SGK1^20^, cortisol also binds to MR under similar conditions^21,22^. It subsequently induces SGK1 expression in human epithelial cells by activating glucocorticoid response elements.^23^ There is a strong correlation between high-salt intake, SSBP, and mutations in the glucocorticoid receptor (GR) gene *NR3C1,* whose product regulates many mammalian genes, such as those for inflammatory cytokines^24,25^. However, the upstream mechanisms regulating SGK1-ENaC activation in APCs leading to SSBP are not known. Here, we characterized the role of cortisol and the APC GR-dependent regulatory role of SGK1 in SSBP.

## METHODS

### Human Studies

Following approval from the Institutional Review Board of Vanderbilt University Medical Center for all experimental procedures, we adhered strictly to the principles outlined in the Declaration of Helsinki and required written consent prior to enrollment. We studied two cohorts described previously^7,8^.

### Phenotyping of SSBP: Salt-Loading and Salt-Depletion Protocol

#### -Study Population

In cohort 1, twenty-five hypertensive participants ages 18–65, with systolic blood pressure (SBP) >130 mmHg or diastolic blood pressure (DBP) >80 mmHg were recruited at Vanderbilt University Medical Center (VUMC) from 2019–2023. In cohort 2, we conducted RNA sequencing on human monocytes after high-salt treatment *in vitro*. The monocytes were isolated from eleven healthy women aged between 22-46 years. Individuals were excluded for 1) renovascular or endocrine causes of secondary hypertension, 2) infectious or inflammatory disease (i.e., active infection or connective tissue disorder), 3) a history of an acute cardiovascular event within six months of the study, 4) treatment-induced high blood pressure or (e.g., selective serotonin reuptake inhibitors and serotonin and norepinephrine reuptake inhibitors, chronic use of decongestants or non-steroidal anti-inflammatory drugs), 5) treatments that alter the immune response (e.g., immunomodulators, immunosuppressants, glucocorticoids), or 6) pregnancy. Demographic and clinical data were collected from the participants and their clinical charts. A physical exam and blood pressure measurements using an automated Omron HEM-907XL monitor were obtained during the screening visit. Plasma and urine EETs were measured via liquid chromatography-mass spectrometry.

#### -Study Protocol

Antihypertensive medications were discontinued two weeks before the study visit for those on antihypertensive treatment. Subjects were instructed to maintain their usual diet and activity level during this period. To ensure the safety of participants who stopped taking their medications, we instructed patients to check their blood pressure twice a day, while seated, after resting for 5 minutes, and to inform the study physician of the results. One day before admission, subjects were instructed to collect a 24-hour baseline urine sample. Participants were admitted to the VUMC Clinical Research Center for three nights to assess salt sensitivity using an inpatient salt loading and depletion protocol, as described previously in detail^26^. The participants were continuously monitored by the study physician. A normal dinner was provided to the participants on admission day. An ambulatory blood pressure monitor (Spacelabs 90207) was placed on the participants the next morning. Baseline blood samples were drawn at 8 AM before any intervention. Salt loading (on day 1) was achieved with a diet containing 160 mEq NaCl (prepared by the University of Alabama Bionutrition Core of the Clinical Research Unit Metabolic Kitchen) and with a 2 L intravenous infusion of normal saline, administered from 8 AM to 12 PM. A 24-hour urine sample was collected from 8 AM on day 1 to 8 AM on day 2. Participants were advised to retire at 10 PM each day. The effects of salt depletion were measured on day 2. Blood samples were collected at 8 AM after overnight fasting to measure salt loading. Salt was depleted using three 40-mg doses of oral furosemide administration at 8 AM, 12 PM, and 4 PM and a low-salt diet containing 10 mEq NaCl. Two 12-hour urine samples were collected from 8 AM to 8 PM to measure the effects of furosemide and from 8 PM on day 2 to 8 AM on day 3 to measure salt depletion. The third blood sample was drawn at 8 AM on day 3 to measure salt depletion. The participants were discharged following blood and urine collection. BP and pulse rate were recorded every 15 minutes from 6 AM to 10 PM and every 30 minutes at night throughout the three study days. The average of the blood pressure recordings from 6 AM to 8 AM on day 1 was used as baseline blood pressure. The blood pressures recorded between 12 PM and 10 PM on days 1 and 2 were used to calculate the average blood pressures during salt loading and depletion, respectively. No specific threshold was used to define SS or SR, but rather, salt sensitivity was analyzed as a continuous variable.

Throughout the study, participants had unlimited access to water; however, their food intake was limited to the diet provided according to the protocol. Before interventions, body weight was recorded at baseline and daily at 7 AM. Body mass index (BMI) was calculated as weight in kilograms divided by height in square meters.

### Laboratory Analysis

Laboratory data including electrolytes and creatinine, plasma renin concentration, and aldosterone were analyzed at the VUMC Pathology Laboratory. Plasma renin direct renin/renin mass and plasma aldosterone were measured using chemiluminescent radioimmunoassay. Blood was collected in EDTA tubes, centrifuged at room temperature, and separated immediately. Urine specimens for four periods (24-hour baseline, 24-hour salt loading, 12-hour furosemide-induced diuresis, and 12-hour salt depletion) were collected and refrigerated as described previously^27^.

### Cellular Indexing of Transcriptomes and Epitopes by Sequencing (CITE-Seq)

As previously described^8^, we performed cell hashing and CITE-Seq analysis on PBMCs of nine participants from cohort 1, depicted in Table 1 (Cohort1). Blood samples were collected at baseline (day 1), after salt load (day 2), and salt depletion (day 3), as shown in Figure 1A. PBMCs from each day were used for cell hashing and CITE-Seq analysis. Per the manufacturer’s protocol (Fisher Scientific, Cat# 14-959-51D), we used BD Vacutainer^®^ CPT^TM^ Mononuclear Cell Preparation tubes to isolate PBMCs. We used antibody-oligonucleotide conjugates to stain the isolated PBMCs and performed sample multiplexing with sample-specific hashtags. We introduced the mixture of cells and an antibody-oligonucleotide complex into a microfluidic system to capture the unique homing guide RNA (hgRNA) barcodes using custom beads containing poly-dT and specific nucleotide sequences. The VANTAGE core generated small vesicles containing a single bead and a single cell through oil inclusion, which encapsulated the antibody-oligonucleotide complex. Then, the cell was lysed, followed by the synthesis of cDNA and antibody-derived tags through reverse transcription.

**Figure 1:**
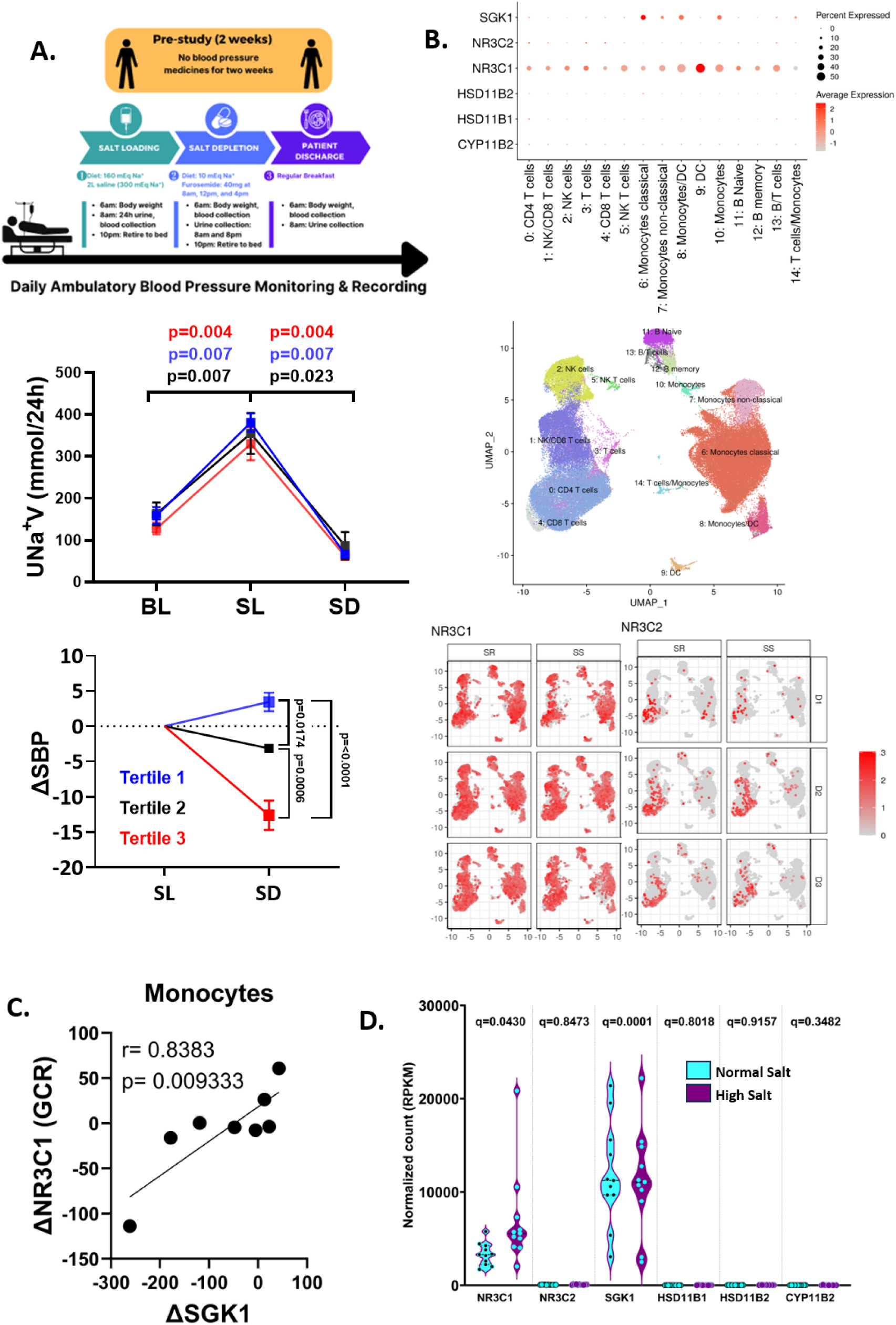
Salt-sensitivity of Blood Pressure Phenotyping and NR3C1 and SGK1 gene expression. **A)** Diagram of the 72-hour inpatient salt-loading (SL) and salt-depletion (SD) protocol with changes in urinary sodium excretion (UNa+V) and systolic blood pressure (ΔSBP). **B)** Dot plot and Uniform Manifold Approximation and Projection (UMAP) representations of relevant gene expression in different immune cell type clusters after CITE-Seq analysis. Patients with the highest salt-sensitivity index were indicated as salt sensitive while those with the lowest were indicated as salt resistant. Days 1, 2, and 3 represent baseline, salt loading, and salt depletion, respectively. **C)** Correlation between changes in gene expression of *SGK1* and *NR3C1* from SL to SD. **D)** Violin plots comparing gene transcript counts via bulk RNA-Seq after normal-salt and high-salt treatments (n=11). Adjusted p values for each pairwise comparison are shown using false discovery rate (FDR < 0.05). NR3C1 indicates Nuclear Receptor Subfamily 3, Group C, Member 1; NR3C2, Nuclear Receptor Subfamily 3, Group C, Member 2; SR, salt-resistant; SS, salt-sensitive.

**Table 1.**
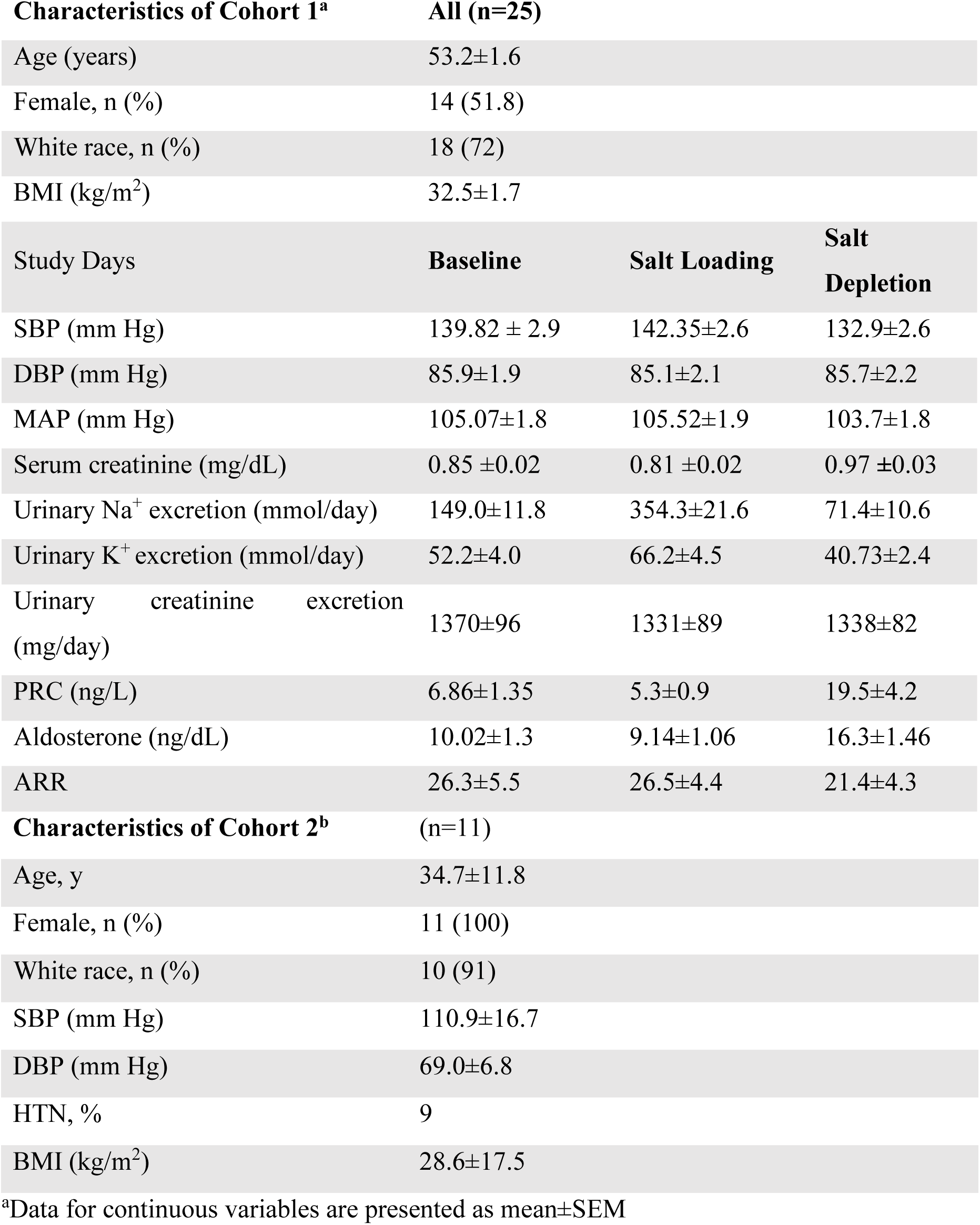
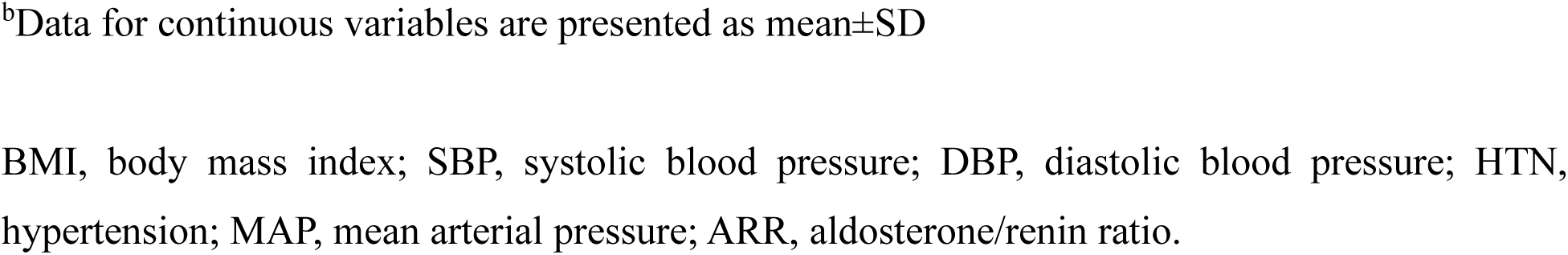
Demographic and clinical characteristics of the participants used in Figure 1D (Cohort 2) along with those assessed for SSBP (Cohort 1).

| **Characteristics of Cohort 1<sup>a</sup>** | **All (n=25)** |  |  |
| --- | --- | --- | --- |
| Age (years) | 53.2±1.6 |  |  |
| Female, n (%) | 14 (51.8) |  |  |
| White race, n (%) | 18 (72) |  |  |
| BMI (kg/m<sup>2</sup>) | 32.5±1.7 |  |  |
| **Study Days** | **Baseline** | **Salt Loading** | **Salt Depletion** |
| SBP (mm Hg) | 139.82 ± 2.9 | 142.35±2.6 | 132.9±2.6 |
| DBP (mm Hg) | 85.9±1.9 | 85.1±2.1 | 85.7±2.2 |
| MAP (mm Hg) | 105.07±1.8 | 105.52±1.9 | 103.7±1.8 |
| Serum creatinine (mg/dL) | 0.85 ±0.02 | 0.81 ±0.02 | 0.97 ±0.03 |
| Urinary Na<sup>+</sup> excretion (mmol/day) | 149.0±11.8 | 354.3±21.6 | 71.4±10.6 |
| Urinary K<sup>+</sup> excretion (mmol/day) | 52.2±4.0 | 66.2±4.5 | 40.73±2.4 |
| Urinary creatinine excretion (mg/day) | 1370±96 | 1331±89 | 1338±82 |
| PRC (ng/L) | 6.86±1.35 | 5.3±0.9 | 19.5±4.2 |
| Aldosterone (ng/dL) | 10.02±1.3 | 9.14±1.06 | 16.3±1.46 |
| ARR | 26.3±5.5 | 26.5±4.4 | 21.4±4.3 |
| **Characteristics of Cohort 2<sup>b</sup>** | **(n=11)** |  |  |
| Age, y | 34.7±11.8 |  |  |
| Female, n (%) | 11 (100) |  |  |
| White race, n (%) | 10 (91) |  |  |
| SBP (mm Hg) | 110.9±16.7 |  |  |
| DBP (mm Hg) | 69.0±6.8 |  |  |
| HTN, % | 9 |  |  |
| BMI (kg/m<sup>2</sup>) | 28.6±17.5 |  |  |

<sup>a</sup>Data for continuous variables are presented as mean±SEM
<sup>b</sup>Data for continuous variables are presented as mean±SD
BMI, body mass index; SBP, systolic blood pressure; DBP, diastolic blood pressure; HTN, hypertension; MAP, mean arterial pressure; ARR, aldosterone/renin ratio.

We used 10x Genomics Cell Ranger 6.0.2 equipped with in-house scripts to quantify genes and demultiplex sample-specific hashtags. The hashtag abundance cutoff for positive cells was determined by the modified R package cutoff. Every cell was categorized as 1) a singlet with a specific hashtag, 2) a doublet, or 3) negative. The genotypic results obtained from Souporcell were combined with the results based on hashtags.^28^ Subsequently, clustering analysis was performed using Seurat^28^ set to a resolution of 1.0. The cell type of each cluster was classified based on the cell activity database^29,30^ and manually refined according to cell-specific marker genes. The program edgeR^31^ was used to identify differential gene expression across conditions with absolute fold change larger than 1.5 and FDR adjusted p value less than 0.05. The WebGestaltR^32^ package was used for Genome Ontology and KEGG pathway over-representation analysis on differentially expressed genes. GSEA^33^ was used for gene enrichment analysis.

### Bulk RNA Sequencing and Bioinformatics

As described previously,^34^ we isolated peripheral blood mononuclear cells (PBMCs) from 40 mL of heparinized blood from eleven healthy subjects and depicted in Table 1 (Cohort 2). Using a Ficoll-gradient protocol, monocyte isolation kit (Miltenyi Biotec 130-091-151) was used to extract monocytes via magnetic labeling and negative selection. Monocytes were cultured in 12-well plates at a density of 1 × 10^6^/mL in Roswell Park Memorial Institute media 1640 (Gibco) supplemented with 10% fetal bovine serum, 1% penicillin/streptomycin, 1% HEPES, and 0.05 mM 2-mercaptoethanol with either 150 mM or 190 mM Na^+^. Bulk RNA sequencing of isolated monocytes grown in a normal or high-salt medium *in vitro* was done by the Vanderbilt Technologies for Advanced Genomics (VANTAGE) core to ensure a high RNA integrity number. The VANTAGE core used an Illumina Tru-Seq RNA sample prep kit to perform polyadenylated RNA sequencing, and paired-end sequencing was done on the Illumina HiSeq2500. FASTQ data from the paired-end sequencing analysis were aligned against the human GRCh38 reference genome assembly with TopHat 2 using the R package. The quaternary round of quality control for raw data and alignment was conducted using QC3, and the MultiRankSeq method was used for expression analysis. A false discovery rate (FDR < 0.05) was used to correct for multiple hypothesis testing.

### Cortisol Analysis

The metabolite contents of the urine and plasma samples from the 72-hour protocol of cohort 1 (**Figure 1A**, **Table 1**) were extracted from the global untargeted metabolomic analysis by Metabolon which uses ultra-high-performance liquid chromatography/tandem accurate mass spectrometry. Raw data for each biochemical detected on a per-sample basis was expressed as peak area (i.e., integrated area-under-the-curve), and these raw values were used in correlation analysis.

### Statistical Analysis

Statistical analysis was performed with Prism version 10.2 (GraphPad Software, La Jolla, USA). Statistical significance was set to a p-value of 0.05. Correlation analyses were performed to compare multiple interval or ratio variables. Associations of continuous variables of interest were assessed with Spearman’s rank correlation and Pearson’s correlation test, as specified in the respective figure legends. Trend lines and confidence intervals (CI) were estimated by linear regression.

## RESULTS

### The Glucocorticoid Receptor, but not the Mineralocorticoid Receptor is expressed in Myeloid Antigen Presenting Cells

To determine whether activation of APC SGK1 is downstream of the mineralocorticoid or the glucocorticoid receptor, we performed single cell CITE-seq analysis on people assessed for SSBP using a highly rigorous modified Weinberger protocol of salt-loading and salt-depletion in humans illustrated in **Figure 1A** and as previously reported.^3,5^ **Table 1**/**Cohort 1** shows the demographics of the study participants. As we have previously published, we found that the blood pressure responses of participants to salt loading/depletion could be categorized into three main tertiles^14^. Tertile 1 (blue) included the inversely salt-sensitive participants, Tertile 2 included the SR participants, and Tertile 3 (red) included the most salt-sensitive participants. In contrast to these very different changes in blood pressure, all subjects show virtually identical changes in 24-hour urinary Na^+^ excretion during salt loading and depletion. These results show that renal handling of Na^+^ is not different between SS and SR people and suggest a role for extra-renal mechanisms in SSBP. As shown in **Figure 1B**, we found that while SGK1 and the GR (indicated by NR3C1) are highly expressed in the APCs—including the monocytes and dendritic cells—there is little to no expression of the MR (indicated by NR3C2) in these cells. **Figure 1B** shows the Uniform Manifold Approximation and Projection (UMAP) to sub-clustered immune cell types, and how the expression of NR3C1 and NCR3C2 changes from baseline (D1) to salt-loading (D2) and salt-depletion (D3) in SR and SS people. Accordingly, we found that changes in the expression of SGK1 from salt loading to salt depletion correlated with changes in NR3C1 (**Figure 1C**). To confirm these findings, we performed bulk RNA-sequencing on isolated monocytes from humans treated with either normal-sodium (140 mM) or high-sodium concentrations (190 mM) in vitro. As shown in **Figures 1B and 1D**, we found significant expression of NR3C1 and SGK1 but little to no expression of NR3C2. Moreover, in accordance with our previous publication, immune cells exhibited minimal to no expression of aldosterone synthase, encoded by CYP11B2, and both 11β-HSDs, encoded by HSD11B1 and HSD11B2.^35^ Since these enzymes are necessary for the synthesis of aldosterone and the conversion of cortisol and cortisone, our results suggest that aldosterone and cortisol are not synthesized in immune cells and point to an important role for the glucocorticoid receptor in APC SGK1-ENaC activation in SSBP.

### Urine and Plasma Cortisol/Cortisone Correlate with Salt Sensitivity of Blood Pressure

To determine if cortisol plays a role in SSBP, we measured cortisol and its precursor cortisone in the plasma and urine of people phenotyped for SSBP. We employed two delta variables in the correlation analysis to investigate the effect of overall salt balance on blood pressure measures and cortisol metabolism. Specifically, we analyzed the differences between salt loading and baseline blood pressure measures, cortisol, and cortisone values to represent the effect of salt loading on blood pressure and cortisol metabolism. Additionally, we included the difference between salt loading and salt depletion to investigate the acute response to salt depletion. This approach allows for a comprehensive analysis of the effects of salt balance on blood pressure and cortisol metabolism. We found that changes in urine cortisol positively correlated with changes in SBP and PP but not DBP or MAP (**Figure 2A**). We found no significant correlations between changes in plasma cortisol with changes in SBP, DBP, MAP or PP (**Figure 2B**). Changes in urine cortisone were not significantly correlated with changes in SBP, DBP, MAP or PP (**Figures 2C**). We found a positive correlation between changes in plasma cortisone with changes in SBP and MAP but not with DBP or PP (**Figure 2D**). These results suggest a role for urine and plasma levels of cortisol and cortisone respectively in SSBP.

**Figure 2:**
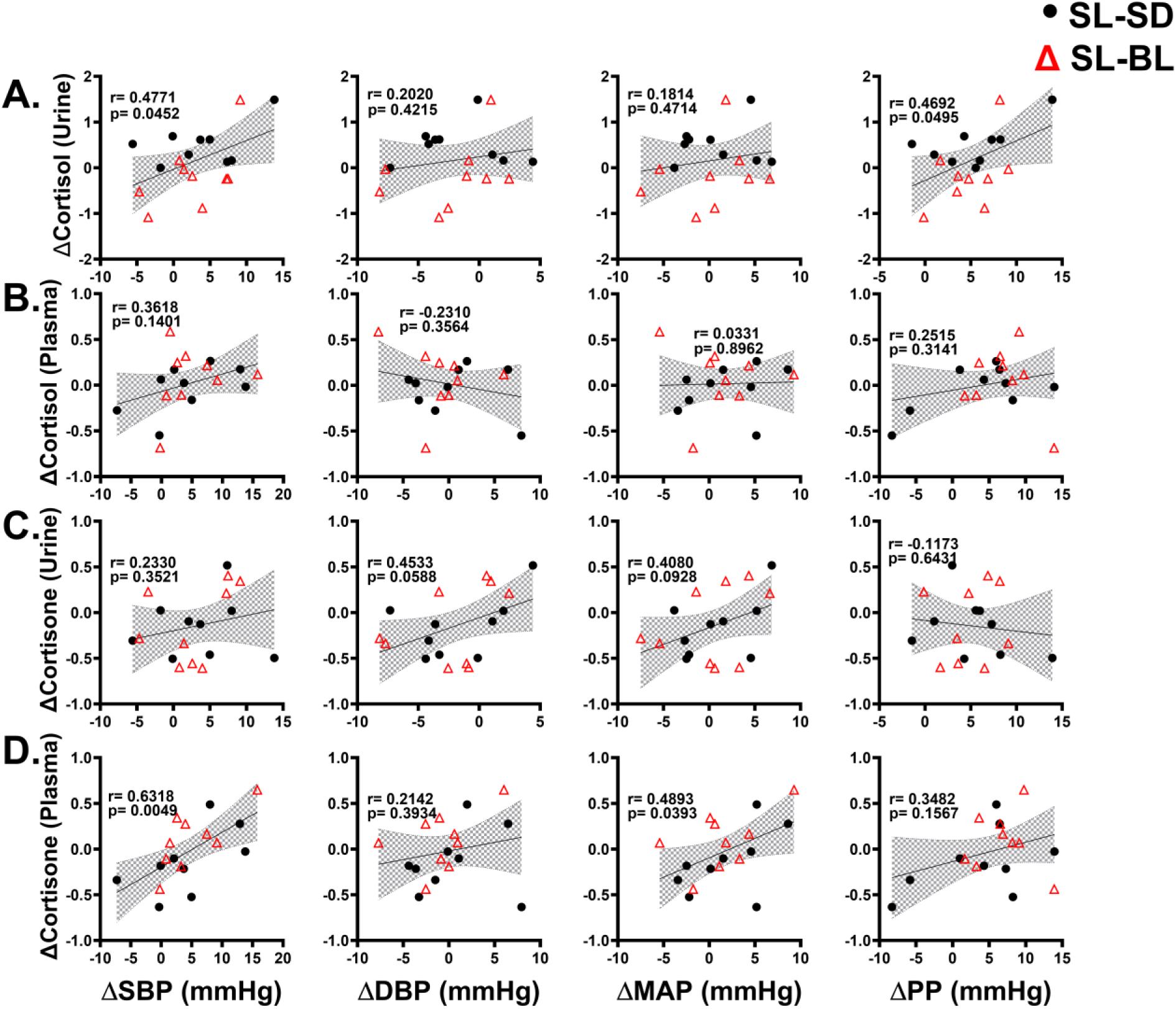
Glucocorticoid metabolome influences hemodynamics. **(A-D)** Scatter plots demonstrating the correlations between blood pressure and changes in **(A)** urine cortisol, **(B)** plasma cortisol, **(C)** urine cortisone, and **(D)** plasma cortisone from mass-spectroscopy analysis. Each metabolite is measured for changes in systolic blood pressure (ΔSBP), diastolic blood pressure (ΔDBP), mean arterial pressure (ΔMAP), and pulse pressure (ΔPP). SL-SD represents difference between the values of salt loading and salt depletion, while SL-BL represents the difference between the values of salt loading and baseline. Trend lines and confidence intervals were estimated using linear regression. The significance and r values were computed using Pearson’s correlation test. Normality of distribution was assessed using Shapiro-Wilk test.

### Urine Cortisol Positively Correlates with Myeloid Cell activation via IsoLGs and negatively associates with Epoxyeicosatrienoic Acids (EETs)

To determine whether cortisol activates immune cells via IsoLG formation, we performed flow cytometry to analyze IsoLGs present in APCs using a gating strategy shown in **Figure 3A**. We found that urine cortisol levels positively correlated with IsoLG accumulation in classical monocytes, non-classical monocytes, and intermediate monocytes. (**Figure 3B**). In addition, we found that plasma EET8-9, EET11-12, EET14-15 and total EETs negatively correlated with urine cortisol. Whereas urine EET8-9 and EET11-12 did not correlate, EET14-15 and total EETs significantly correlated with baseline urine cortisol (**Figure 3C**). These results suggest that EETs and cortisol play a role in APC ENaC-mediated IsoLG production in SSBP.

**Figure 3:**
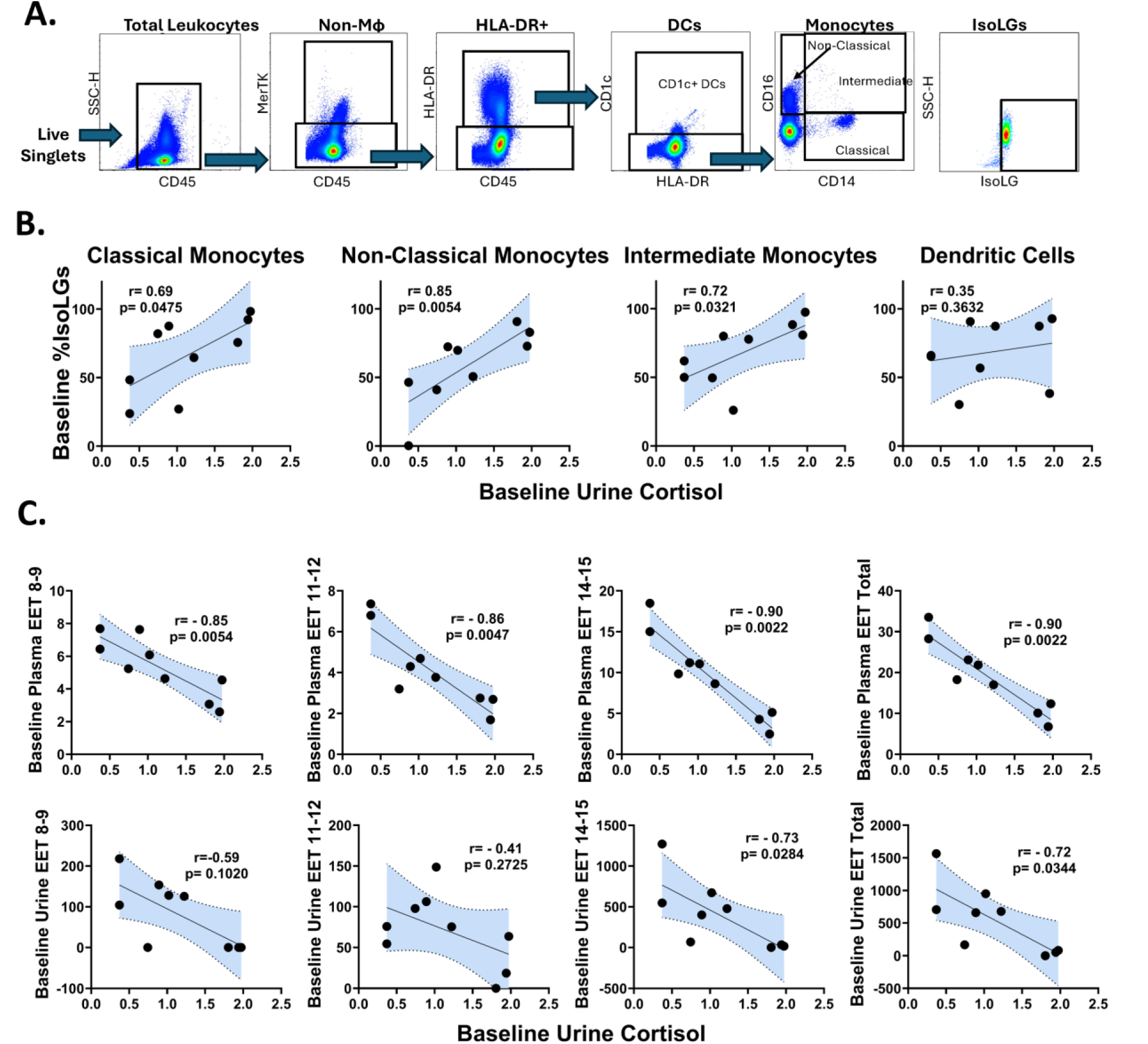
Urine cortisol is positively correlated with IsoLG percentage and negatively correlated with plasma EETs. **(A)** Gating strategy to identify IsoLG-adducts in dendritic cells (DCs) and classical, intermediate, and non-classical monocytes. **(B)** Scatter plots demonstrating the correlations between baseline percent of IsoLGs and baseline urine cortisol in classical monocytes, non-classical monocytes, intermediate monocytes, and dendritic cells. **(C)** Correlations of baseline plasma and urine EETs with baseline urine cortisol in classical monocytes, non-classical monocytes, intermediate monocytes, and dendritic cells. Trend lines and confidence intervals were estimated with linear regression and the significance and r value were computed using Spearman’s rank correlation.

### Aldosterone and renin are not associated with immune cell activation via IsoLGs in SSBP

To determine the role of aldosterone and renin in immune cell activation via IsoLGs in SSBP, we measured APC IsoLGs using flow cytometry, as well as aldosterone, renin, and the aldosterone-renin ratio in plasma. We found no correlation of baseline APC IsoLGs with aldosterone, renin, and aldosterone-renin ratio (**Figure 4A**). Moreover, we did not find any significant correlations between changes in APC IsoLGs and aldosterone, renin, or aldosterone-renin ratio (**Figure 4B**).

**Figure 4:**
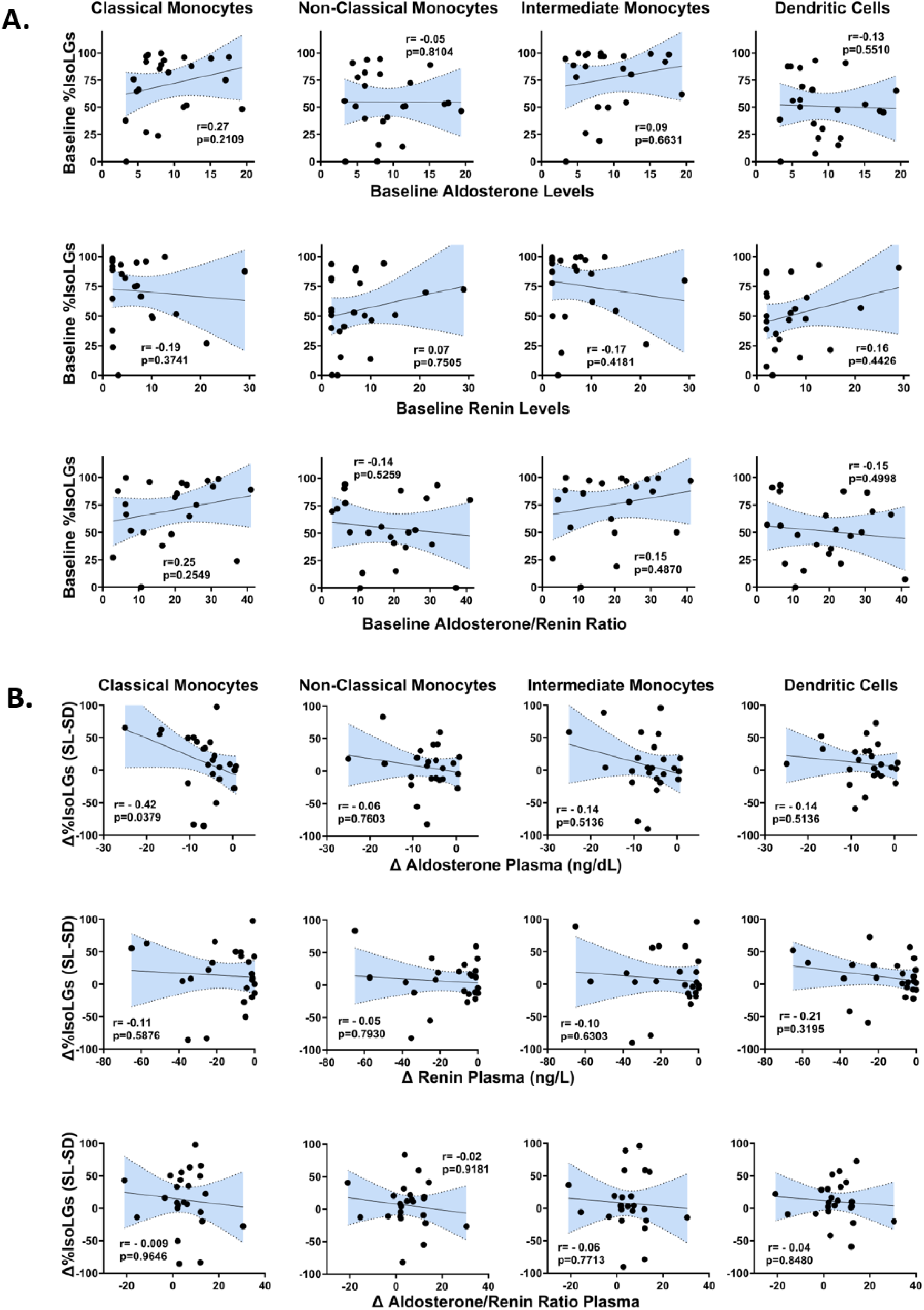
Correlations of IsoLGs with aldosterone and renin. **(A)** Scatter plots demonstrating the correlations between baseline percent of IsoLGs and baseline aldosterone, renin, and aldosterone-renin ratio in classical monocytes, non-classical monocytes, intermediate monocytes, and dendritic cells. **(B)** Correlations of changes in percent of IsoLGs with changes in plasma aldosterone, renin, and aldosterone-renin ratio in classical monocytes, non-classical monocytes, intermediate monocytes, and dendritic cells. All the delta (Δ) variables were calculated as the difference between values on salt loading and salt depletion days (SL-SD). Trend lines and confidence intervals were estimated with linear regression. R value and the significance were computed using Spearman’s rank correlation.

## DISCUSSION

We recently found that human immune cells express a unique sodium channel composed of ENaCα and ENaCδ, which is not found in the human kidney, and its expression patterns change with blood pressure following salt loading and salt-depletion in SSBP.^8,15,36^ We also previously found that salt-induced expression of ENaC in immune cells is regulated by SGK1^15^ and that changes in SGK1 expression in APCs parallel blood pressure changes during salt-loading/depletion in people with SSBP^36^. Data presented in the current study indicate that the GR, not the MR, signaling via cortisol is responsible for myeloid APC ENaC regulation upstream of SGK1 (**Figure 5**). Our studies show that myeloid APCs do not express the MR but rather express the GR which was associated with expression of SGK1 in these cells. In this study, we also found that cortisol, not aldosterone, is associated with SSBP and immune cell activation via IsoLGs. Implication of the GR-SGK1-ENaC signaling pathway in SSBP underscore the unique pathogenesis of this independent risk factor for CVD and the importance of developing unique immune targeted treatments different from salt-resistant hypertension.

**Figure 5:**
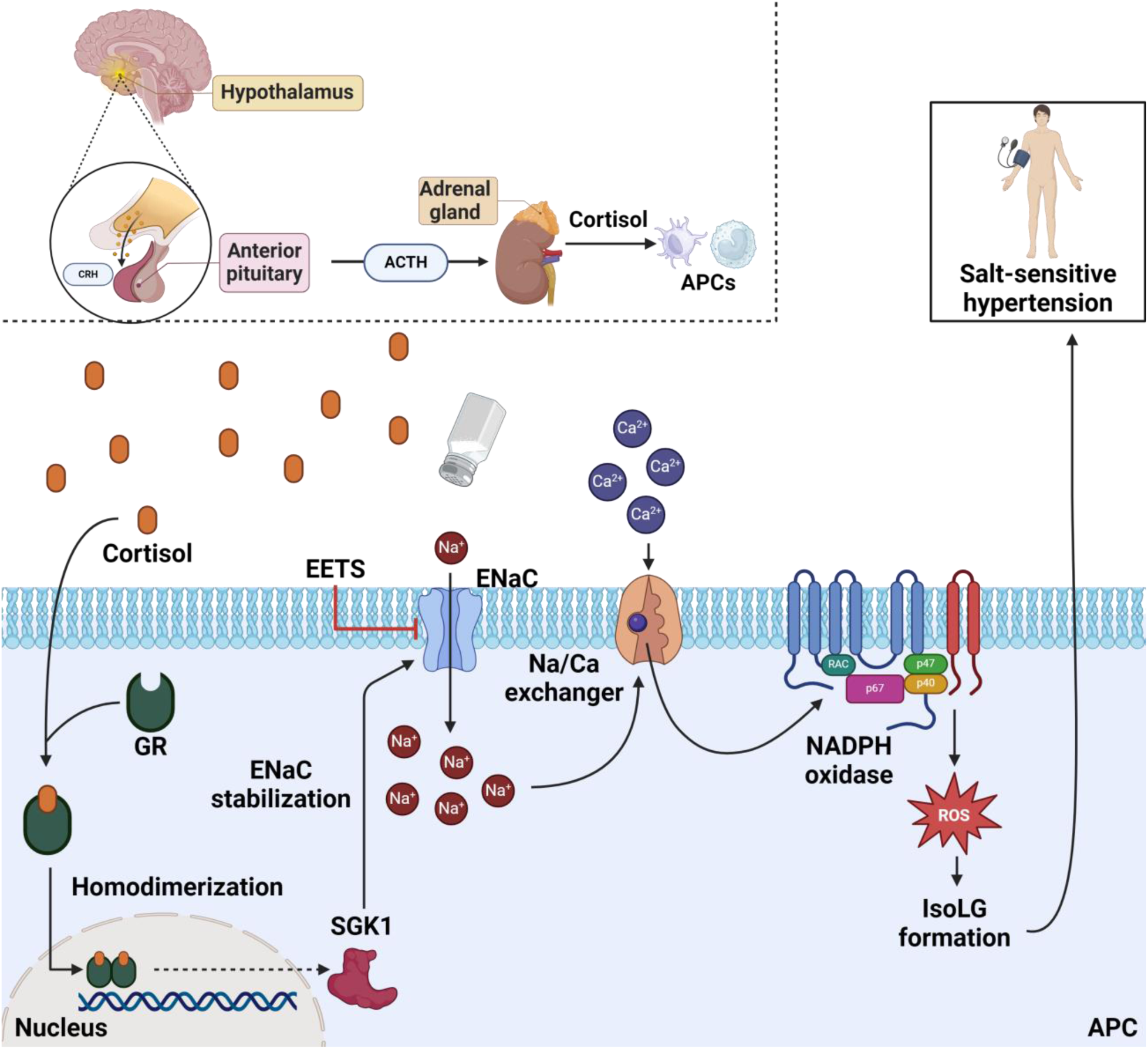
Proposed mechanism for the role of cortisol in SSBP. The release of cortisol from the adrenal gland is a direct response to adrenocorticotropic hormone (ACTH) production by the anterior pituitary gland, which is controlled by the hypothalamus. Cortisol activates glucocorticoid receptor, likely inducing *SGK1* expression for the regulation of epithelial sodium channel (ENaC) in monocytes. High levels of extracellular sodium enter the monocytes via ENaC to cause an influx of calcium ions through the Na^+^/Ca^2+^ exchanger. Calcium induces NADPH oxidase activation, producing reactive oxygen species (ROS) and IsoLGs. Subsequent T-cell activation and renal damage are events that are indicative of salt-sensitive hypertension.

Previous studies have shown that Blacks have a higher prevalence of SSBP than Whites, yet they have lower plasma levels of aldosterone, enhanced response to amiloride and paradoxical increases in response to salt with higher expression as well as sensitivity of SGK1.^37,38^ Moreover, mice lacking SGK1 are protected against salt-sensitive hypertension induced by a high-fat diet.^39^

We demonstrated previously the essential role of APC SGK1 in SSBP.^8,15^ Whereas the role of SGK1 activation downstream of MR activation by aldosterone has been extensively studied in chronic salt-induced hypertension secondary to renal perturbations,^40^ the mechanism of SGK1 activation in the context of acute blood pressure changes according to salt-intake (SSBP) has not been studied. Here we show that myeloid APCs lack MR expression but instead, we demonstrate high levels of GR expression in these cells (**Figure 1**).

In previous studies, we showed that chronic treatment of salt and aldosterone induces blood pressure elevations secondary to renal tubular remodeling.^40^ However, we found that plasma aldosterone levels do not correlate with SSBP in humans.^35^ In the current studies, we demonstrate that changes in both plasma and urine cortisol correlate with human SSBP (**Figure 2**). Moreover, cortisol but not aldosterone correlates with immune cell activation via IsoLGs (**Figures 3 and 4**).

Loss-of-function mutations in *HSD11B2* are strongly associated with SSBP, and cortisol excess often correlates with hypertension^41^. Glucocorticoids, mineralocorticoids, and inflammatory factors stimulate the production of SGK1^42^. Although GR, also known as *NR3C1*, is expressed widely in immune cells, its expression is highest in monocytes, as is *SGK1* (**Figure 1D**). Typically, the GR forms a nuclear transcription factor complex to attenuate inflammatory responses ^43^. We found that independent of MR, *NR3C1* gene expression increased in hypertensive patients after treatment with high salt, suggesting an alternative proinflammatory mechanism of high-salt-dependent SGK1 activation by GR. Our research indicates that SGK1 regulates ENaC-dependent sodium entry into APCs, leading to SSBP and inflammatory response through IsoLG formation by the NRPL3 inflammasome and IL-1β production.^8^

We also showed that myeloid cell ENaC-induced inflammation and SSBP are regulated independently of the renin-angiotensin-aldosterone system (RAAS).^35^ Here, we provide further evidence of this, as we observed no correlation between baseline levels of aldosterone, renin, and the aldosterone/renin ratio (ARR) with %IsoLG levels in immune cells (**Figure 4A**). Moreover, the differences in %IsoLG levels between salt loading and depletion did not correlate with any of these measures (**Figure 4B**). These results suggest the regulation of immune cell ENaC in SSBP through the GR pathway rather than the RAAS system.

It is worth noting that the prevalence of SSBP is higher in women than in men across all ethnicities and increases with age in both genders, raising the question of whether a cortisol-dependent pathway plays a role in sex and age related differences in SSBP.^44,45^ Previous research has indicated that cortisol levels are higher in men who are salt-sensitive compared to those who are salt-resistant^46^. However, to date, there are limited studies that investigate gender differences in cortisol metabolism. One review suggested that loss of estrogen increases the risk for SSBP in postmenopausal women^45^. While the intricate effect of estrogen on SSBP is not well known currently, the increase in SSBP prevalence in menopause can be attributed to age related increase in cortisol alone, as increased cortisol levels are also associated with age and late-stage menopause.^47,48^. It should also be noted that psychosocial stressors affect cortisol metabolism and diurnal response of cortisol levels and linked to poorer health outcomes which may also affect social, racial and gender differences in SSBP.^49^

An important aspect to consider is the interpretation of plasma cortisol levels, as they reflect the pulsatile nature of the hypothalamus-pituitary-adrenal (HPA) axis. This pulsatile release occurs in approximately 90-minute intervals and peaks in the early morning, which coincides with our plasma sampling period. Conversely, urine cortisol levels are sustained over a longer time period.^50^ Indeed, urine cortisol levels are typically considered a more reliable measurement and are frequently used as a diagnostic tool for conditions involving glucocorticoid excess, such as Cushing’s syndrome. Collectively, this suggests that urine cortisol would serve as a more reliable diagnostic biomarker for SSBP when compared to plasma cortisol

Our study has several limitations. First, the limited sample size did not permit us to investigate whether the difference in cortisol excretion in urine could explain the observed race and gender differences in SSBP. Furthermore, the design of our study and the limited availability of the human cells make it difficult to draw direct causal inferences regarding GR-induced SGK1 expression and IsoLG production in APCs. Instead, it allows us to explore the existence of such an association and further mechanistic studies with larger sample sizes are necessary to confirm such differences and causality. Despite these limitations, our single-cell non-supervised evidence with confirmation of bulk RNA studies that myeloid APC do not express the MR, but instead express the GR are highly rigorous. Moreover, to our knowledge, this is the first study to demonstrate that GR, not MR, responds to dynamic changes in dietary salt intake in humans. This novel finding provides a new direction for further research into the role of immune cortisol-GR signaling in SSBP.

In summary, our results suggest that cortisol signaling contributes to immune cell activation and SSBP rather than aldosterone signaling. These findings indicate a novel glucocorticoid-mediated mechanism for SSBP that may parallel stress-induced hypertension.

## Author Contributions

A.K., T.K., and M.S. conceived and designed the research; C.F.A., M.S., J.A.I., A.L.M. and A.P.H. performed the experiments; M.S., K.N., Q.S., M.D., J.A.I., A.L.M., C.L.L., C.N.W., H.K.B., A.G.M., Z.V., and A.H. analyzed the data; C.F.A., M.D., T.K., and A.K. interpreted the results of the experiments; M.S, Q.S., C.L.L., K.N., A.G.M., A.H.J., H.K.B., M.D., A.L.M., and Z.V. prepared the figures. C.F.A., M.D., and K.N. drafted the manuscript; A.H. and A.K. edited and revised the manuscript; T.A.I, V.K., A.H. and A.K. approved the final version of the manuscript. All authors have reviewed and agreed to the current version of the manuscript.

## SOURCES OF FUNDING

This study was supported by the National Institutes of Health grants D34HP16299, T32AI007281, T32GM14492, T32HL00773 (CFA), R01HL147818 (AK and TK), T32HL144446 (AP), R03HL155041 (AK), R01HL144941 (AK), Doris Duke CSDA 2021193 (CNW), K23 HL156759 (CNW), Burroughs Wellcome Fund 1021480 (CNW), the Vanderbilt CTSA grant UL1TR002243 from NCATS/NIH, NIH grants DK059637 and DK020593 to Vanderbilt University Medical Center Hormone Assay and Analytical Services Core, R01HL138519 (AKH), R56AG068026 (ERG). The UNCF/Bristol-Myers Squibb E.E. Just Faculty Fund, Career Award at the Scientific Interface (CASI Award) from Burroughs Wellcome Fund (BWF) ID # 1021868.01, BWF Ad-hoc Award, NIH Small Research Pilot Subaward to 5R25HL106365-12 from the National Institutes of Health PRIDE Program, DK020593, Vanderbilt Diabetes and Research Training Center (DRTC) Alzheimer’s Disease Pilot & Feasibility Program. CZI Science Diversity Leadership grant number 2022-253529 from the Chan Zuckerberg Initiative DAF, an advised fund of the Silicon Valley Community Foundation (AHJ), AHA career development award 23CDA1053072 (MS).

## Notes

### Competing Interest Statement

The authors have declared no competing interest.

